# Comprehensive analysis of intercellular communication in thermogenic adipose niche

**DOI:** 10.1101/2022.09.28.509990

**Authors:** Farnaz Shamsi, Rongbin Zheng, Li-Lun Ho, Kaifu Chen, Yu-Hua Tseng

**Affiliations:** Section on Integrative Physiology and Metabolism, Joslin Diabetes Center, Harvard Medical School, Boston, MA 02115, USA; Department of Molecular Pathobiology, College of Dentistry, New York University, New York, NY 10010, USA; Department of Cell Biology, Grossman School of Medicine, New York University, New York, NY 10016, USA; Basic and Translational Research Division, Department of Cardiology, Boston Children’s Hospital, Boston, MA 02115, USA; Department of Pediatrics, Harvard Medical School, Boston, MA 02115, USA; Computer Science and Artificial Intelligence Laboratory, Massachusetts Institute of Technology, Cambridge, MA, USA; Harvard Stem Cell Institute, Harvard University, Cambridge, MA, USA

## Abstract

Brown adipose tissue (BAT) is responsible for regulating body temperature through adaptive thermogenesis. The ability of thermogenic adipocytes to dissipate chemical energy as heat counteracts weight gain and has gained considerable attention as a strategy against obesity. BAT undergoes major remodeling in a cold environment. This remodeling results from changes in the number and function of brown adipocytes, expanding the network of blood vessels and sympathetic nerves, and changes in the makeup and function of immune cells. All these processes are essential for enhanced BAT thermogenesis to maintain euthermia in the cold. Such synergistic adaptation requires extensive crosstalk between the individual cells in tissues to coordinate their responses. To understand the mechanisms of intercellular communication in BAT, we applied the CellChat algorithm to single-cell transcriptomic data of mouse BAT. We constructed an integrative network of ligand-receptor interactome in BAT and identified the major signaling input and output of each cell type. By comparing the ligand-receptor interactions in BAT of mice housed at different environmental temperatures, we found that cold exposure enhances the intercellular interactions among the major cell types in BAT, including adipocytes, adipocyte progenitors, lymphatic and vascular endothelial cells, myelinated Schwann cells (MSC), nonmyelinated Schwann cells (NMSC), and immune cells. Furthermore, we identified the ligands and receptors that are regulated at the transcriptional level by temperature. These interactions are predicted to regulate the remodeling of extracellular matrix (ECM), inflammatory response, angiogenesis, and neurite growth. Together, our integrative analysis of intercellular communications in BAT and their dynamic regulation in response to housing temperatures establishes a holistic understanding of the mechanisms involved in BAT thermogenesis. The resources presented in this study provide a valuable platform for future investigations of BAT development and thermogenesis.

## Introduction

BAT is a specialized adipose type primarily responsible for regulating body temperature through adaptive thermogenesis. The ability of thermogenic adipocytes to dissipate chemical energy in the form of heat counteracts energy storage and has gained considerable attention as a strategy against obesity. Studies in rodents have demonstrated that increasing the amount or activity of brown adipocytes promotes energy expenditure, improves insulin sensitivity, and protects animals from diet-induced obesity[1–3].

The conventional view of cold-induced thermogenesis involves the sensation of cold by the cutaneous and deep body sensory neurons, which in turn signals to the hypothalamic networks that integrate them with brain temperature information to drive appropriate sympathetic neural outflow to BAT[4]. Acute cold exposure stimulates adaptive thermogenesis by enhancing the thermogenic function of existing brown adipocytes. In contrast, prolonged cold exposure promotes de novo recruitment of brown adipocytes and simultaneous remodeling of other adipose resident cells to maximize thermogenesis[5]. Specifically, chronic exposure to cold induces a coordinated expansion of brown adipocyte progenitors, endothelial cells, and nerve terminals, as well as changes in the composition of BAT-resident immune cells[6, 7]. Although adipocytes are the heat-producing cells in BAT, other cell types form the adipocyte niche and regulate adipocyte number and function through extensive cellular crosstalk.

In recent years, single-cell RNA-sequencing (scRNA-seq) analyses of different adipose depots have provided a comprehensive map of adipose resident cell populations, including adipocytes and their progenitors, fibroblasts, different types of immune cells, endothelial cells, vascular smooth muscles, and Schwann cells[8–10]. By analyzing the cell-type-specific transcriptional changes in BAT of mice housed at different temperatures, we showed that cold exposure instigates significant transcriptional changes in all BAT-resident cells. Many transcriptional changes are accompanied by cell identity transitions, e.g., the differentiation of Trpv1-expressing progenitors to thermogenic adipocytes[8]. However, how distinct cell types in the BAT microenvironment communicate and how the responses of individual cells are integrated to produce a coordinated adaptation have not been described.

Here, we use scRNA-seq data from BAT of mice housed at different temperatures to construct the ligand-receptor interactome within the BAT microenvironment. Using CellChat to quantitatively infer intercellular crosstalk from the cell-type specific gene expression data, we identify the primary signaling inputs and outputs for each cell type in BAT. This analysis reveals the role of adipocyte progenitors as the major communication “hub” in the adipose niche. Notably, comparing the ligand-receptor interactome of BAT in mice housed at different temperatures revealed that cold exposure significantly enhances cellular crosstalk in BAT. Specifically, the number of incoming and outgoing interactions to or from adipocytes, adipocyte progenitors, lymphatic and vascular endothelial cells, myelinated Schwann cells (MSC), non-myelinated Schwann cells (NMSC), and several immune cell types increases in cold. By comparing the expression of ligands and receptors in each cell type, we identify the interactions that are significantly regulated by ambient temperature. Collectively, our integrative analysis of cellular communication in BAT and their dynamic responses to cold provide new insights into the mechanisms of BAT thermogenesis and cold adaptation. This resource will pave the way and serve as a guide for future investigations of BAT development and function.

## Results

### Housing temperature shapes the cellular composition of BAT

To understand how different cell types in BAT respond to changes in ambient temperature, we isolated the stromal vascular fraction (SVF) of BAT from mice housed at thermoneutrality (TN: 30 °C for 7 days), room temperature (RT: 22 °C) or cold (5 °C for 2 days or 7 days)[8]. We used the total single cell suspensions and profiled their transcriptome using the 10x Genomics Single Cell 3’ Gene Expression solution. The resulting dataset included 107,679 high-quality cells and 468 million reads. Cells were clustered using a graph-based approach (Scanpy)[11] and visualized using Uniform Manifold Approximation and Projection (UMAP) method[12]. Using a combination of known markers and the specific transcripts expressed in each cluster, we assigned cell type identities to the clusters. Overall, we identified 20 distinct clusters representing major cell types in mouse BAT (Figure 1a). We have previously reported the identification and characterization of the non-hematopoietic cells in BAT and their cold-induced adaptations using a fraction of this dataset[8]. In this study, we used the full dataset to comprehensively analyze the effects of housing temperature on cellular composition and intercellular communications in the BAT microenvironment.

**Figure 1.**
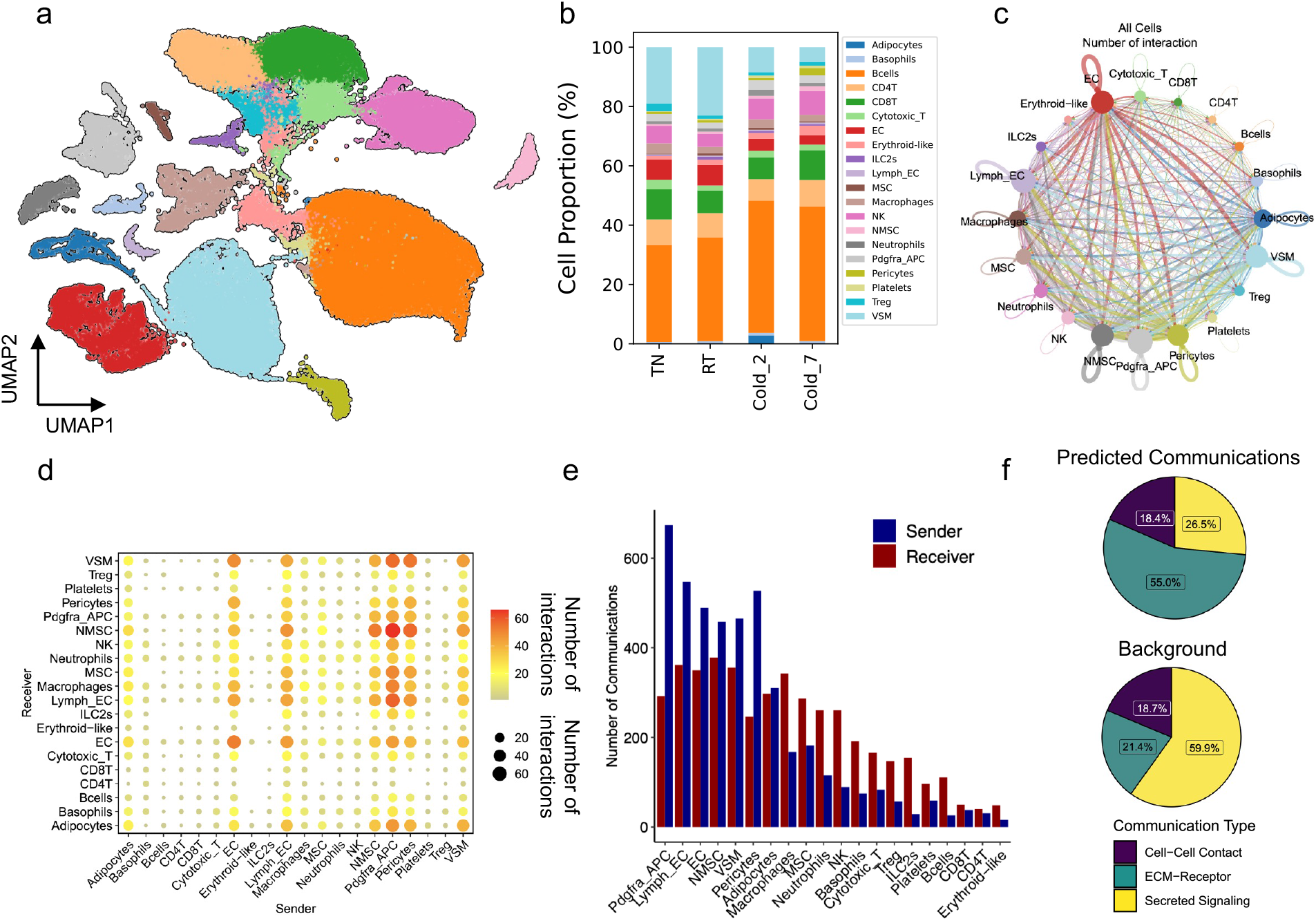
Overview of cellular composition and cell-cell interactions in BAT. (a) Unsupervised clustering of cells from the BAT-SVF of 9-week-old male C57BL/6J mice housed at TN (30 °C, 7 days), RT (22 °C), Cold (5 °C, 2 and 7 days) represented on a UMAP. (b) The percentage of cells in each cluster under different housing conditions. (c) Circle plot showing the cell-to-cell communications in BAT. The line width represents the number of ligand-receptor interactions between two cell types. The size of the circle represents the number of interactions in each cell type, line width represents the number of interactions between two cell types, and color distinguishes sender cell types. (d) Dot plot showing the number of significant interactions between each pair of cell types. The size and color of the circle both represent the number of interactions, respectively. (e) Number of significant communications involving each cell type as “sender” or “receiver”. (f) The proportion of communications in Cell-Cell contact, ECM-Receptor, and Secreted Signaling categories among the predicted communications (top) and background (bottom).

Comparing the cellular composition of BAT from mice housed at different environmental temperatures revealed significant differences at the cell type level between the sample groups (Figure 1b and Supplementary Figure 1a-t). Notably, a significantly higher proportion of adipocytes were detected in mice housed at cold for 2 days (Wald test p-value 0.0005) (Supplementary Figure 1a). Given that the SVF isolation step was used to deplete the lipid-laden adipocytes, these remaining adipocytes likely represent the differentiating adipocytes with lower lipid content and buoyancy than the fully mature adipocytes.

In this analysis, we identified multiple immune cell types within the mouse BAT microenvironment. These include B cells, several populations of T cells, such as CD4+T, CD8+T, cytotoxic T, and regulatory T (Treg) cells, erythroid-like cells, basophils, neutrophils, natural killer (NK) cells, macrophages, and type 2 innate lymphoid cells (ILC2) (Figure 1a and 1b). Interestingly, colder housing temperature increased the proportion of ILC2s (Wald test p-value 0.0029) (Supplementary Figure 1m) and reduced the proportions of cytotoxic T-cells (Wald test p-value 0.0297) (Supplementary Figure 1f) and regulatory T cells (Tregs) (Wald test p-value 0.0026) in BAT (Supplementary Figure 1g). ILC2s are the source of Th2 cytokines IL5 and IL13 that promote the recruitment and accumulation of eosinophils and alternatively activated macrophages[13]. The rise in the proportion of ILC2s in BAT of mice housed in cold is consistent with previous work showing that ILC2s directly regulate thermogenic adipocyte development by regulating the number and fate of adipocyte precursors [14, 15]. Previous studies have revealed the contribution of adipose-resident T cells to adipose function and systemic metabolism[16–18]. The pro-inflammatory cytotoxic T cells were shown to be upregulated in the epididymal white adipose tissue (WAT) of diet-induced obese mice and contribute to the propagation of adipose inflammation[19]. Although B cells constituted the largest immune cell population within the BAT niche, their proportions were not affected by housing temperature.

Additionally, mice housed at colder ambient temperatures showed larger proportions of lymphatic ECs (Wald test p=0.0278) (Supplementary Figure 1p), non-myelinating Schwann cells (Wald test p=0.0041) (Supplementary Figure 1t), pericytes (Wald test p=0. 0466) (Supplementary Figure 1r), and lower proportions of vascular smooth muscles (Wald test p=0.0305) in BAT (Supplementary Figure 1q). This is an intriguing observation, especially considering the recent identification of Trpv1-expressing cells derived from the vascular smooth muscles (VSM) as the source of cold-induced thermogenic adipocytes[8]. The changes in the proportions of various vascular cell populations also reflect the dynamic remodeling of BAT vasculatures in response to environmental temperature changes.

### Quantitative inference of the intercellular communications in BAT microenvironment

Intercellular communication enables the individual cells in tissues to coordinate their functions and orchestrate development, homeostasis, and remodeling. Beyond cataloging the cellular composition of tissues, the cell type-specific transcriptional signature can be used to probe the intercellular interactions mediated by secreted and membrane-bound proteins[20]. To map the intercellular crosstalk in BAT mediated by protein-ligand and receptor interactions, we used the CellChat algorithm[21]. CellChat uses a database of more than 2000 interactions among ligands, receptors, and their cofactors representing the known heteromeric molecular complexes to infer the potential communications between cell types in scRNA-seq data. CellChat takes cell-type specific gene expression data as input and computes the likelihood of cell-cell interaction by integrating gene expression with prior knowledge of the interactions between known ligands and receptor complexes. The main advantage of CellChat over other similar methods is that it considers the composition of ligand-receptor complexes, including multimeric ligand and receptor complexes and cofactors such as soluble agonists, antagonists, co-stimulatory and co-inhibitory membrane-bound receptors.

CellChat identified a total of 4438 significant ligand-receptor pairs among the 20 cell groups. We used these interactions to construct the network of intercellular communications among all cell types (Figure 1c-d). Constructing a weighted directed graph of significant connections between the cell types indicated that Pdgfra+ adipocyte progenitor cells (APC) were involved in the most significant number of interactions with other cell types (Figure 1c and 1e). Pdgfra+ adipocyte progenitors send and receive signals via 674 and 292 ligand-receptor pairs, respectively. This indicates the role of adipocyte progenitors as the dominant communication “hub” in the adipose niche and suggests the multifaceted roles of adipocyte progenitors in the adipose microenvironment beyond supporting adipogenesis. Vascular cells including lymphatic and vascular endothelial cells, vascular smooth muscles, and pericytes are also engaged in a large number of communications with other cell types in the adipose niche (Figure 1c and 1e).

Classifying the intercellular communications revealed that most crosstalk in BAT is mediated by the interactions of cells with extracellular matrix (ECM) proteins (55% of the identified communications) while signaling through the secreted molecules and cell-cell contact make up 26.5% and 18.4% of the communications, respectively (Figure 1f). This distribution is different from the background dataset in which secreted signaling, ECM-receptor interactions, and cell-cell contact represent 59.9%, 21.4%, and 18.7% of interactions, respectively (Figure 1f). This is in agreement with the crucial roles of ECM components in the regulation of adipose tissue function and remodeling[22, 23] and suggests that the interaction of cells with the surrounding ECM in BAT is an important regulator of their functions.

### Cold exposure promotes intercellular communications in BAT

To determine how ambient temperature affects intercellular communications in BAT, we applied CellChat to analyze cell-cell interactions in each group (temperature) separately. As environmental temperatures decreased, the total number of interactions increased. Cold exposure significantly increased the number of interactions in BAT (3586 for TN, 3246 for RT, 4251 for cold2, 5281 for cold7, p-value < 2.2e-16) (Figure 2a-b). This was reflected in significant increases in the number of both incoming and outgoing interactions from and to several cell populations. For example, 7 days of cold exposure increased the number of incoming and outgoing interactions to and from the Pdgfra+ adipocyte progenitors, lymphatic and vascular endothelial cells, non-myelinated Schwann cells (NMSC), and myelinated Schwann cells (MSC), adipocytes, and several immune cell types (Macrophages, neutrophils, natural killer cell, ILC2s, etc.). Interestingly, the analysis also indicated that the Pdgfra+ APC received the highest number of incoming signals after 7 days of cold. At that time, NMSC, VSM, and EC were the cell types that sent out significant outgoing signals.

**Figure 2.**
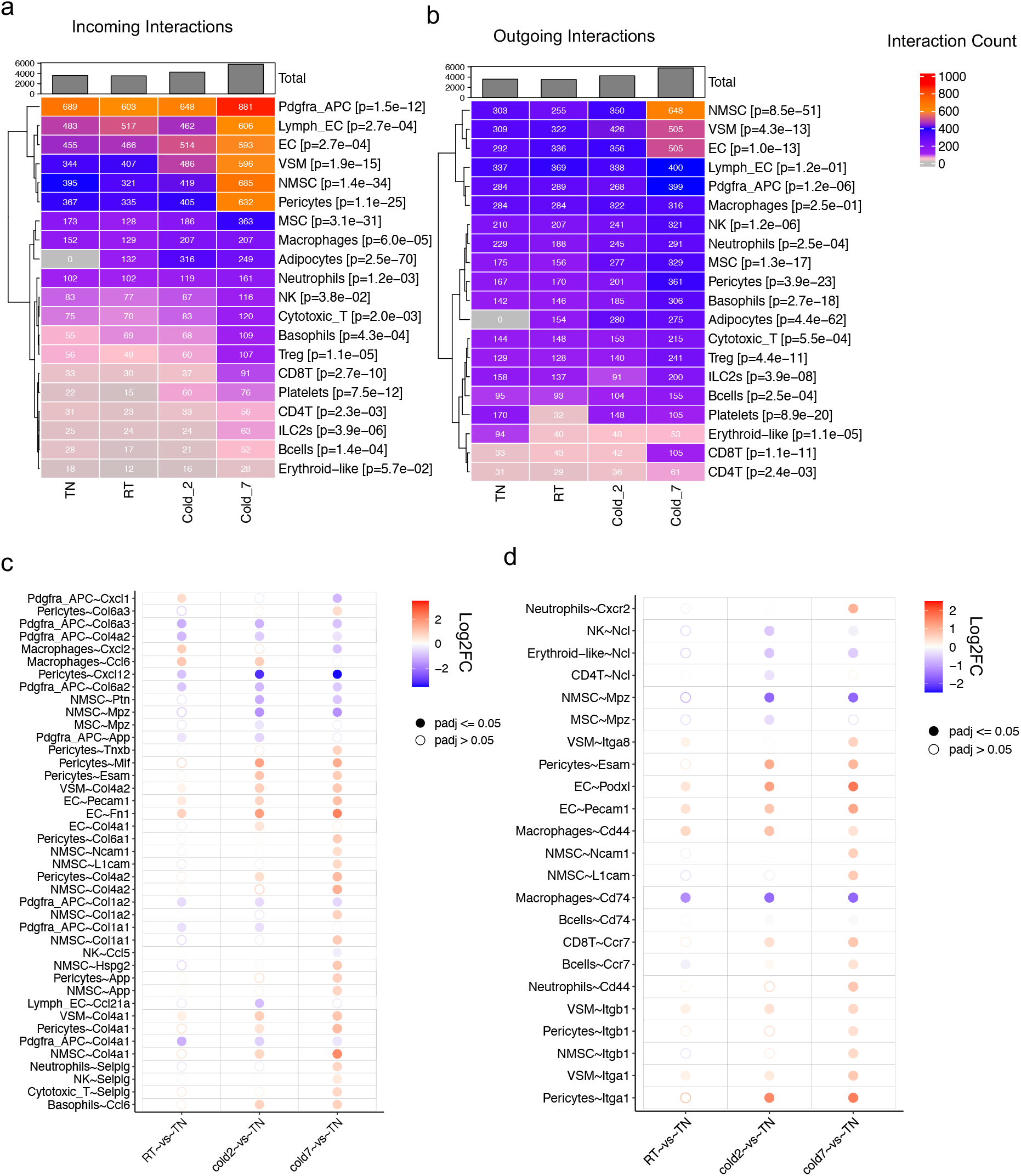
Temperature-regulated remodeling of intercellular interactions in BAT. Heatmaps showing the total number of significant (a) incoming and (b) outgoing interactions for each cell type across different conditions. The expression fold change of (c) ligands and (d) receptors relative to the TN (30 °C, 7 days) group. The color indicates log2 fold change of gene expression between two conditions. The solid dots indicate significant differential expression (p-value < 0.05) and circle dots indicate non-significance.

Cold exposure stimulates a coordinated and multilayered remodeling of BAT, including the recruitment of brown adipocytes, expansion of vascular endothelial cells, sympathetic nerve outgrowth, and changes in the makeup of BAT resident immune cells[5, 8, 24]. The profound increase in the number of interactions among different cell types in the BAT indicates the higher need for effective communications between cells to coordinate their functions in response to thermogenic stimuli.

### Temperature-regulated remodeling of intercellular interactions in BAT

Next, we analyzed the effect of housing temperature on the expression of individual ligands and receptors mediating crosstalk in BAT. Using K-means clustering on communication probabilities of 125 most variable communication events across conditions, we identified five distinct patterns for temperature-dependent regulation of interactions (Supplementary Figure 2a). Among those, interactions that follow Pattern 1 (upregulated in 7 days cold vs. other groups), Pattern 2 (upregulated with decreasing temperature across all groups), and Pattern 4 (downregulated with decreasing temperature across all groups) are most likely to be involved in adaptive BAT thermogenesis. The 46, 31, and 24 interactions representing patterns # 1, 2, and 4 are presented in Supplementary Figure 2b, respectively.

Analyzing the expression of individual ligands revealed that cold exposure regulates the RNA expression level of cytokines or chemokines (Ccl6, Ccl21a, Ccl5, Mif, Cxcl12, Cxcl2, Cxcl1), ECM proteins (Col4a1, Col1a1, Col1a2, Col4a2, Col6a1, Col6a2, Col6a3, Hspg2, Fn1, Tnxb, Ptn), and cell adhesion proteins (Selplg, L1cam, Ncam1, Pecam1, Esam1, Mpz) in immune cells, vascular cells, and Schwann cells (Figure 2c). Furthermore, cold exposure induced changes in the expression of multiple receptors on adipose resident cells (Figure 2d). For example, cold exposure significantly increased the expression of Cd44 and Cxcr2 in neutrophils, while it reduced the expression of Cd74 in macrophages (Figure 2d). Other significantly regulated ligands and receptors in the vascular cells include the upregulation of Esam and Col4a1/2 in pericytes, Itga1 and Itgb1 in vascular smooth muscles, and Pecam1 and Fn1 in vascular endothelial cells (Figure 2c-d). Collectively, these changes can shift how these niche components interact with each other to enable the functional and structural adaptation required for BAT thermogenesis.

### Crosstalk between adipogenic and immune cells

Adipose tissue depots house a wide array of immune cells, including macrophages, B and T lymphocytes, NK cells, neutrophils, eosinophils, basophils, and ILC2s. Emerging evidence revealed several roles for the resident immune cells in adipose thermogenesis. Our scRNA-seq data identified several distinct populations of immune cells in BAT and enabled us to analyze the effect of housing temperature on the abundance and transcriptome of each population. To understand the ligand-receptor crosstalk between the immune cells and adipogenic cells, we mapped the interactions between the individual immune cell populations present in the scRNA-seq data (Basophils, B-cells, CD4 T-cells, CD8-T cells, Cytotoxic T-cells, ILC2s, Macrophages, Neutrophils, NK cells, and Tregs) and adipogenic cells (adipocytes, Pdgfra+ APC, and VSM) (Figure 3a). Notably, the Pdgfra+ APCs acted as major sender cells to communicate with macrophages, NK cells, and neutrophils. Among all the interactions between adipogenic and immune cells, 43% were between ECM and receptors, 32.9% were through secreted factors, and 24.1% were mediated via cell-cell contact (Figure 3b).

**Figure 3.**
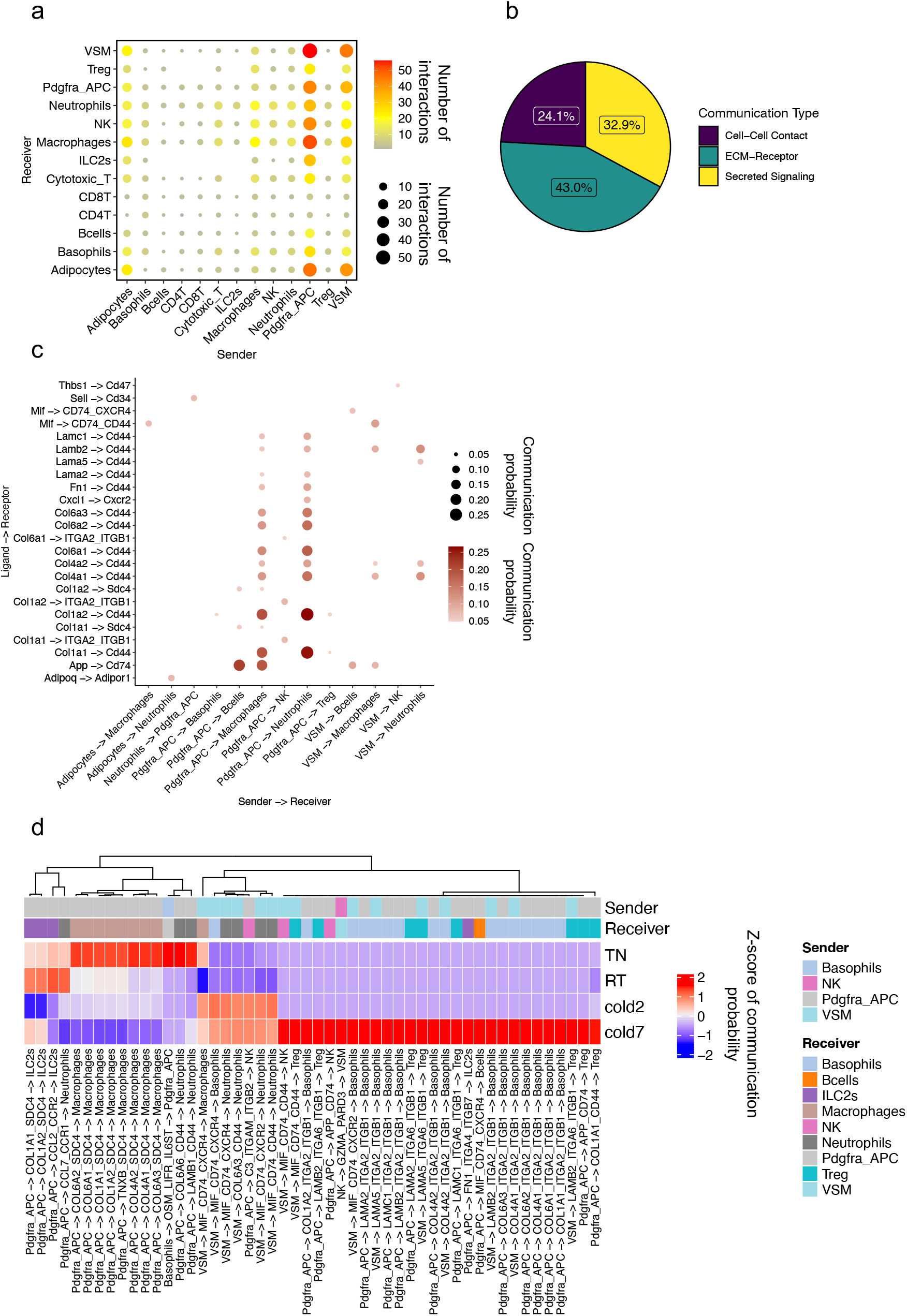
Crosstalk between adipogenic and immune cells. (a) Dot plot showing the number of significant interactions between each pair of cell types. The size and color of the circle both represent the number of interactions. (b) The proportion of communications in Cell-Cell contact, ECM-Receptor, and Secreted Signaling categories among the predicted communications. (c) Dot plot presenting the most significant ligand-receptor interaction pairs between adipogenic and immune cells in BAT. The size and color of the circle both represent communication probability. (d) Heat map showing the communication scores for each ligand-receptor interaction in the cell type pairs across different conditions.

This analysis identified many interactions between the Pdgfra+ adipocyte progenitors and several immune cells. For example, Pdgfra+ adipocyte progenitors secreted amyloid beta (A4) precursor protein (App) that can bind to Cd74 on macrophages (Figure 3c). Stimulation of CD74 is shown to trigger the activation of several signal transduction cascades that play important roles in cell proliferation and survival[25]. Additionally, Pdgfra+ adipocyte progenitors secreted multiple types of collagens, such as Col1a1, Col1a2, Col4a1, Col4a1, Col6a1, Col6a2, and Col6a3, all of which could interact with Cd44 on macrophages or neutrophils (Figure 3c). Cd44 is a transmembrane glycoprotein expressed in a variety of cell types. Cd44 interacts with extracellular matrix components and regulates the migration and recruitment of leukocytes[26].

Differential expression analysis across the four conditions (TN, RT, cold 2 days, and cold 7 days) identified several ligands and receptors whose expression was modulated by cold, resulting in a significant decrease or increase in the interactions as defined by the calculated communication scores. For example, cold exposure significantly reduced the expression of transcripts encoding the ECM components (Col6a2, Col6a1, Col6a3, Col1a1, and Col1a2) in Pdgfra+ adipocyte progenitors and its interacting partner, syndecan 4 (Sdc4) in macrophages (Figure 3d). Furthermore, the reduced expression of C-C motif chemokine ligand 2 (Ccl2) in Pdgfra+ adipocyte progenitors could attenuate the proinflammatory Ccl2-Ccr2 signaling axis in Ccr2 expressing ILC2s in cold (Figure 3d).

Conversely, the interactions between the Macrophage migration inhibitory factor (MIF) ligand expressed by the vascular smooth muscles and the receptor complexes Cd74-Cxcr4, Cd74-Cxcr2, and Cd74-Cd44 expressed on macrophages, basophils, neutrophils, NK, and Treg cells were predicted to increase in BAT of mice housed in cold (Figure 3d). Mif is a pleiotropic cytokine that regulates leukocyte recruitment and plays a role in innate and adaptive immunity[27]. Another example is the increased interactions of the Pdgfra+ adipocyte progenitors with basophils and Tregs via laminins (Lama2, Lamc1, and Lamb2) and integrins (Itga2, Itgb1) (Figure 3d).

### Crosstalk between adipogenic and vascular cells

BAT is one of the most vascularized tissues in the body[28]. Environmental challenges, such as ambient cold, excess calorie intake, and physical activity, promote the expansion of adipose tissue vasculature primarily through inducing sprouting angiogenesis[29]. The expansion and remodeling of the vascular network are essential for providing the optimal supply of oxygen, nutrients, and other bioactive molecules for adipocytes. To understand the reciprocal communications between the cells that form adipose vasculature and adipogenic cells, we next focused on the interactions between vascular cells (vascular endothelial cells, lymphatic endothelial cells, vascular smooth muscle, and pericytes) and adipogenic cells (adipocytes, Pdgfra+ adipocyte progenitors, and vascular smooth muscles) (Figure 4a). Interestingly, more than half (62%) of the identified interactions were between ECM and their receptors, 25% were through secreted factors, and 13% occurred via cell-cell contact (Figure 4b).

**Figure 4.**
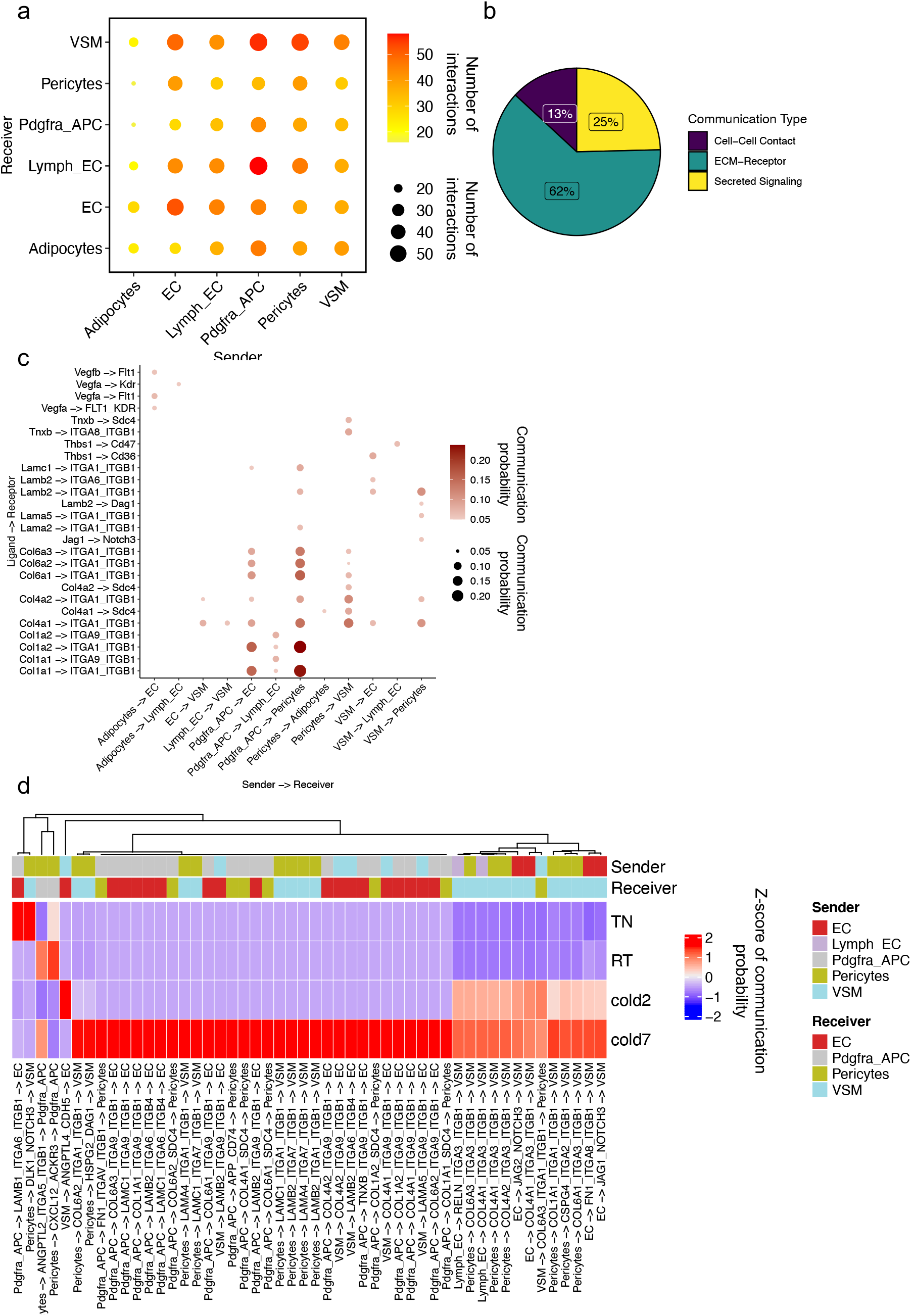
Crosstalk between adipogenic and vascular cells. (a) Dot plot showing the number of significant interactions between each pair of cell types. The size and color of the circle both represent the number of interactions. (b) The proportion of communications in Cell-Cell contact, ECM-Receptor, and Secreted Signaling categories among the predicted communications. (c) Dot plot presenting the most significant ligand-receptor interaction pairs between adipogenic and vascular cells in BAT. The size and color of the circle both represent communication probability. (d) Heat map showing the communication scores for each ligand-receptor interaction in the cell type pairs across different conditions.

Adipocytes secrete angiogenic factors, including members of the vascular endothelial growth factor (VEGF) family that regulate angiogenesis through two tyrosine kinase receptors, VEGFR1 (encoded by *Flt1*) and VEGFR2 (encoded by *Kdr*)[29]. Cellchat identified significant and specific crosstalk between adipocytes and endothelial cells mediated by the interaction of Vegfa and Vegfb with Flt1, Kdr, and the Flt1-Kdr receptor complex (Figure 4c). Pdgfra+ adipocyte progenitors engaged in multiple crosstalks with vascular and lymphatic endothelial cells and pericytes. Most of these interactions appeared to occur through collagens and integrins (such as α1β1 and α9β1) expressed in the vasculature (Figure 4c).

As expected, cold exposure robustly enhanced the interactions between adipogenic and vascular cells (Figure 4d). Cold exposure significantly increased the expression of *Itga9* and *Itgb1* in vascular endothelial cells, potentially resulting in a higher abundance of Integrin α9/β1 complex in cold. This was predicted to enhance their interactions with Pdgfra+ adipocyte progenitors and vascular smooth muscles through the ECM components such as Col6a3, Lamc1, Col1a1, Col6a1, Lamb2, Col4a1, Col4a2, Lama4, and Col6a2.

Cold exposure also induced the expression of Notch ligands, Jag1 and Jag2, in endothelial cells and Notch3 in vascular smooth muscles, thus increasing the likelihood of Jag1/Jag2-Notch3 signaling between the vascular endothelial cells and vascular smooth muscles in cold. The Jag-Notch signaling controls vascular development in embryonic and adult tissues. Specifically, Endothelial cell-derived Jag1 is essential for blood vessel formation by enhancing the differentiation and maturation of vascular smooth muscle cells[30, 31].

### Crosstalk between adipogenic and Schwann cells

BAT is innervated by an extensive network of sympathetic and sensory nerve projections that are responsible for transmitting information between BAT and the central nervous system (CNS)[32]. Cold exposure increases sympathetic activity in BAT by elevating the rate of norepinephrine turnover and increasing the density of sympathetic arborizations[4, 33]. The communication between thermogenic adipocytes and sympathetic nerve plays an essential role in establishing the sympathetic network during the early postnatal period[34].

Although the innervating sympathetic nerves are not captured in the single cell transcriptome analysis, we identified two distinct populations of myelinating and non-myelinating Schwann cells (NMSC and MSC) in BAT (Figure 1a). Schwann cells are the major glial cells of the peripheral nervous system and are essential for the development, function, and regeneration of peripheral nerves [35]. Consistent with their roles in supporting axon growth, we found a significant increase in the frequency of NMSCs in BAT of mice housed in cold (Supplementary Figure 1n). Analysis of ligand-receptor interactions identified many interactions between adipogenic cells (adipocytes, Pdgfra+ adipocyte progenitors, and vascular smooth muscles) and Schwann cells (NMSC and MSC) (Figure 5a). Most of these communications were mediated by ECM-receptor interactions (73%). Signaling by secreted ligand and direct cell-cell contact made up 21% and 6% of total interactions, respectively (Figure 5b).

**Figure 5.**
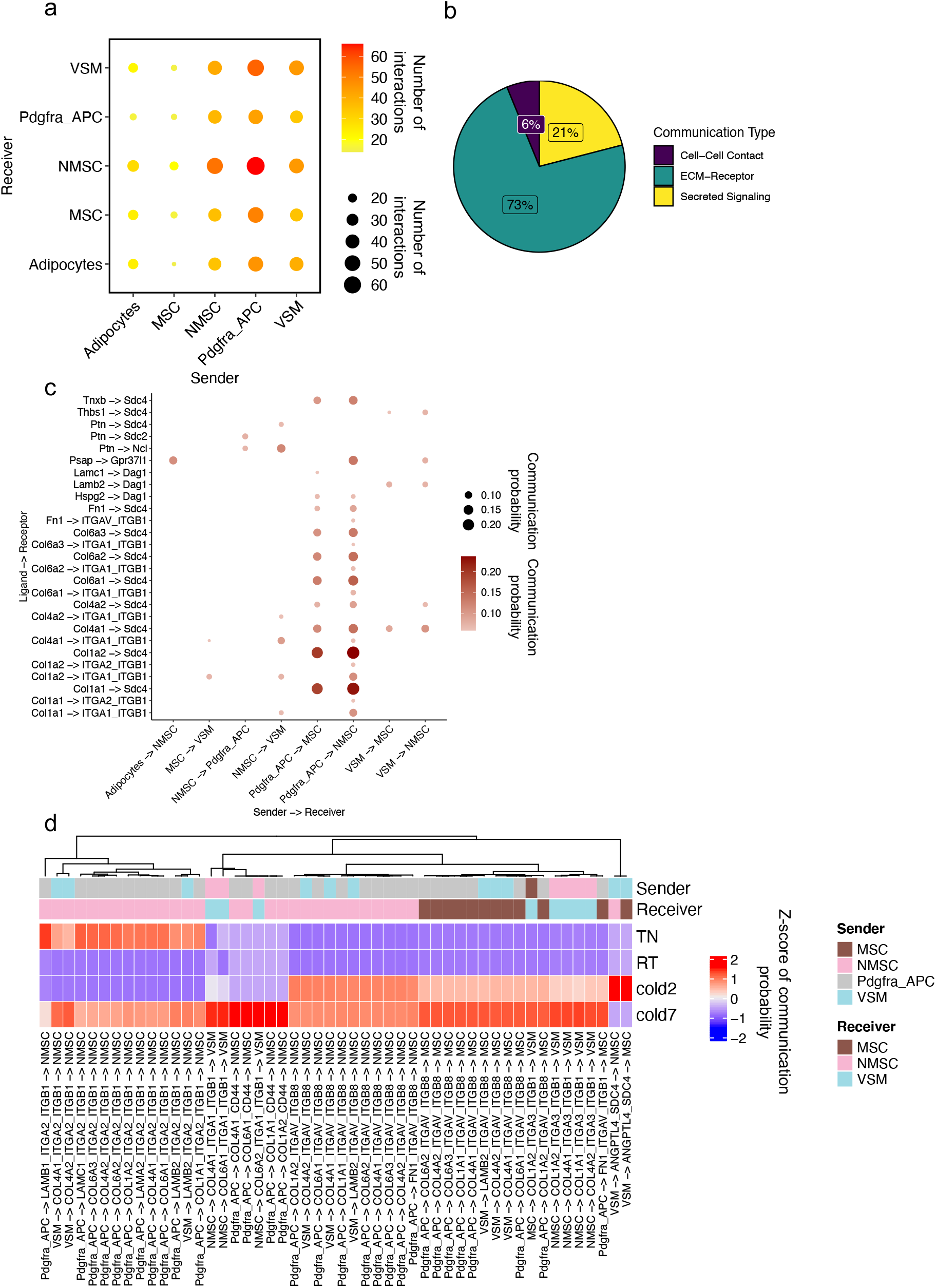
Crosstalk between adipogenic and Schwann cells. (a) Dot plot showing the number of significant interactions between each pair of cell types. The size and color of the circle both represent the number of interactions. (b) The proportion of communications in Cell-Cell contact, ECM-Receptor, and Secreted Signaling categories among the predicted communications. (c) Dot plot presenting the most significant ligand-receptor interaction pairs between adipogenic and Schwann cells in BAT. The size and color of the circle both represent communication probability. (d) Heat map showing the communication scores for each ligand-receptor interaction in the cell type pairs across different conditions.

The largest interactions in this category were between Pdgfra+ adipocyte progenitors and NMSCs. Examples of such interactions included the binding of several ECM proteins expressed in Pdgfra+ adipocyte progenitors (Col1a1, Col1a2, Col4a1, Col6a1, Col6a2, Col6a3, and Tnxb) to syndecan 4 (Sdc4) expressed by MSCs and NMSCs (Figure 5c). Additionally, adipocytes and APCs expressed Prosaposin (Psap) that binds to Gpr37l1 present on NMSCs (Figure 5c). Psap exhibits neurotrophic and myelinotrophic activities in neurons and glial cells [36, 37].

Among the interactions that are differentially regulated by temperature, most are through integrin receptor complexes: α2β1 and α5β8 on NMSCs, α5β8 on MSCs, α1β1 and α3β1 on vascular smooth muscles (Figure 5d). Cold exposure increased the expression of Itga1, Itga3, and Itgb1 in vascular smooth muscles, likely facilitating their interaction with Pdgfra+ adipocyte progenitors through Col4a1, Col6a1, Col6a2, Col1a2, Col4a1, Col1a1, and Col4a2 (Figure 5d).

## Discussion

Applying the CellChat method to single-cell transcriptomic data of mouse BAT, we present a comprehensive map of ligand-receptor interactions in the thermogenic adipose tissue across four different housing temperatures. Our analysis reveals that BAT adaptation to housing temperature involves major changes in the cellular composition as well as the rewiring of the intercellular crosstalk. The quantitative comparison of the ligand-receptor interactions across different temperatures showed that cold exposure promotes intercellular communications in BAT, suggesting the increased need for information exchange among cells to coordinate the tissue adaptation to the rise in energetic demand. This is in line with a recent study showing that obesity and HFD increased the intensity of ligand-receptor pair expression in the white adipose tissue of humans and mouse[38, 39]. These findings highlight the role of intercellular communications in adipose tissue remodeling in response to fluctuating nutritional status and metabolic demand.

Despite great advancement in the characterization of adipocyte progenitors and their contribution to the expansion and turnover of the adipocyte pool, their roles beyond adipogenesis remains poorly understood. Our systematic analysis of ligand-receptor interactions identified adipocyte progenitors as the dominant communication hub in the thermogenic adipose niche. Comparing the transcriptome of adipocyte progenitors across different temperatures demonstrates that housing temperature regulates the transcription of ligands and receptors involved in ECM remodeling, immune modulation, angiogenesis, and neurogenesis. These findings underscore the diverse regulatory functions of adipocyte progenitors in adipose tissue beyond adipogenesis.

Collectively, our integrative analysis of intercellular crosstalk in BAT provides a holistic understanding of the mechanisms that control thermogenesis. The comprehensive database of ligand-receptor interactions identified in this study offers a valuable platform for future investigations of the functions of specific signaling molecules and cell types in BAT development and function.

## Methods

### Animals

All animal experiments were performed in compliance with all relevant ethical regulations applied to the use of small rodents and with approval by the Institutional Animal Care and Use Committees (IACUC) at Joslin Diabetes Center. 9-week-old male C57BL/6J mice (Stock no. 000664) were purchased from The Jackson Laboratory and were used for the scRNA-sequencing experiment.

Mice were housed at room temperature (22°C), cold (5°C for 2 days and 7 days), or thermoneutral (30°C for 7 days) in controlled environmental diurnal chambers (Caron Products & Services Inc., Marietta, OH) with free access to food and water. Mice were maintained at a 12hr-light/dark cycle with 30% Humidity on a normal chow diet containing 22% of calories from fat, 23% from protein, and 55% from carbohydrates (Mouse Diet 9F 5020; PharmaServ). The procedures for the isolation of the stromal vascular fraction from BAT and single cell RNA-sequencing were previously described[8] and outlined below.

### Isolation of the stromal vascular fraction from BAT

Interscapular BAT was dissected, minced, and digested with a cocktail containing type 1 Collagenase (1.5 mg/ml; Worthington Biochemical), Dispase II (2.5 U/mL; STEMCELL Technologies), fatty acid-free bovine serum albumin (2%; Gemini Bio Products) in Hanks’ balanced salt’s solution (Corning^®^ Hank’s Balanced Salt Solution, 1X with calcium and magnesium) for 45 minutes at 37°C with gentle shaking. Dissociated tissue was centrifuged at 500 g at 4°C for 10 minutes. The top adipocyte layer and the supernatant were gently removed, the SVF pellet was resuspended in 10 ml of 10% Fetal Bovine Serum (FBS) in DMEM, filtered through a 100 μm cell strainer into a fresh 50 ml tube, and centrifuged at 500 g for 7 minutes. The red blood cell lysis was performed by resuspending the pellet in 2 ml sterile ACK (Ammonium-Chloride-Potassium) lysis buffer (ACK Lysing Buffer, Lonza) and incubating on ice for 5 minutes. The cells were then filtered through a 40 μm cell strainer, washed with 20 ml 10% FBS in DMEM, and centrifuged at 500 g for 7 minutes. The pellet was resuspended in 1 ml of 1.5% BSA in PBS. The dead cell removal was performed using Dead Cell Removal Kit (Miltenyi Biotec) according to the manufacturer’s instructions. The cells were finally resuspended in 50-100 μl of 1.5% BSA in PBS and kept on ice before immediately proceeding to single cell isolation. The detailed protocol for the isolation of the SVF from adipose tissue is deposited on protocols.io (dx.doi.org/10.17504/protocols.io.bpurmnv6).

### Single-cell RNA-sequencing

Cells were loaded for an expected recovery of 10,000 cells per channel. The chip loaded with single cell suspension was placed on a 10x Genomics Chromium Controller Instrument (10x Genomics, Pleasanton, CA, USA) to generate single cell droplets containing uniquely barcoded GEMs (Gel Bead-In Emulsions). Single-cell RNA-seq libraries were obtained following the 10x Genomics recommended protocol, using the reagents included in the Chromium Single Cell 3’ v3 Reagent Kit. The libraries were sequenced on the NovaSeq S2 flow cell (Illumina, 100 cycles).

### Version of software and packages

Cellranger 6.1.2, Python 3.8; Python packages include scanpy 1.8.2, pandas 1.4.1, matplotlib 3.5.1, seaborn 0.11.2, jupyter 1.0.0; R 4.1.1, R packages include CellChat 1.1.3, ggplot 3.3.5, ggrepel_0.9.1, ComplexHeatmap_2.8.0.

### Single-cell RNA-seq data analysis

The Cellranger was used to align single-cell RNA sequences to the mouse reference genome (mm10) and obtain the read count for each gene in each cell. The outputs in read count in each gene and each cell from Cellranger were provided to Scanpy for further processing. This processing includes filtering low-quality cells and genes, expression normalization, dimensional reduction, clustering, and visualization of gene expression. Cells were filtered if they had less than 800 or more than 50,000 UMIs and if they had less than 400 or more than 7,500 detected genes. Genes were filtered if they were detected in less than 10 cells. Total read counts in each cell were normalized to 10 thousand, and the normalized value of each gene in each cell was log-transformed to obtain the final normalized gene expression. 10 nearest neighbors and 40 principal components were used in clustering and UMAP analysis. Clustering was done by the Leiden algorithm. The visualization of the clusters was performed by the UMAP method (Figure 1a). The clusters were annotated to cell types by known marker genes of cell types and the top differentially expressed genes in the cluster. The known cell type marker genes were collected from PanglaoDB at https://panglaodb.se/markers.html. Differential expression analysis was also performed by Scanpy for each cell type between any two conditions from TN, RT, cold2, and cold7.

### Cell composition analysis

To investigate the cell type composition in BAT from mice housed at four conditions (TN, RT, cold2, and cold7), we calculated the cell proportion score, namely, the percentages of cells in a cell type over the total cell number in the condition. The cell proportion of each cell type and each condition was visualized by a stacked bar plot in Figure 1b. Further, the proportion of a cell type in each mouse and each condition was also calculated and visualized by boxplot. The Wald-test was used to calculate the p-value to suggest the significance of cell composition change across four conditions by each cell type (Supplementary Figure 1).

### Cell-cell communication inference in BAT by CellChat

The normalized expression matrix and cell type annotation generated from Scanpy were passed to CellChat for ligand-receptor cell-cell communication analysis. To quantitatively infer intercellular communications in BAT, all cells from four conditions (TN, RT, cold2, and cold7) were pooled and grouped by cell types for CellChat analysis. Default parameters were used, except that min.cells was set to 10, which allows to filter out cell types with the total number of cells smaller than 10 in CellChat. Mouse ligand-receptor database without any selection was used in CellChat. Circle plot was used to visualize the number of inferred communications between cell types by the netVisual_circle function in the CellChat package (Figure 1c). One communication was counted by a unique combination of sender cell type, ligand, receptor, and receiver cell type.

### Identification of temperature-regulated cell-cell communications

In this analysis, cells were grouped by cell types and conditions to perform CellChat analysis. To identify highly changed communications between conditions (TN, RT, cold2, and cold7), a communication-by-condition matrix was generated and filled by communication probability scores. In this matrix, any communications with a p-value less than 0.05 in one of the four conditions were included. Next, the mean and variance of communication probability were calculated for each row of the matrix, namely, each communication. The temperature-regulated communications were defined if their mean communication probability was greater than 0.05, and the variance greater than 0.0005. The two cutoffs for mean and variance probability were determined based on their distribution across all the communications, respectively. 126 ligand-receptor communications with the largest variation of communication probability across four conditions were selected and considered as temperature-regulated communications.

To better interpret those temperature-regulated communications, k-means clustering was used to group the identified 126 ligand-receptor communications. Five clusters were determined to represent five different patterns of communication across four conditions.

To visualize the five patterns, communication probabilities of all ligand-receptor interactions in the pattern were averaged by each condition, which resulted in four average communications probabilities corresponding to four conditions in each pattern. Then, the averaged communication probabilities for each pattern were shown in a line graph (Supplementary Figure 2a). Further, the communication probability of each communication in patterns was also shown in a heatmap (Supplementary Figure 2b). In the analysis of temperature-regulated communications, adipocyte was not included, since it was mostly differentiating adipocytes, and few cells were included in the TN, cold2, and cold7 conditions.

## Acknowledgments

This work was supported in part by the US National Institutes of Health (grants R01DK077097, R01DK102898, R01DK122808 (to Y.-H.T.), K01DK125608 (to F.S.), and P30DK036836 (to Joslin Diabetes Center’s Diabetes Research Center) from the National Institute of Diabetes and Digestive and Kidney Diseases, CZF2019-002454 from the Chan Zuckerberg Foundation (to Y.-H.T.), R01GM125632 (to K.C.) from NIGMS, and R01HL133254 (to K.C.) and R01HL148338 (to K.C.) from NHLBI.

## Data availability

The genomic data have been deposited to GEO and are available by the accession number GSE160585.

## Supplementary Figures

**Supplementary Figure 1. Ambient temperature changes the cellular composition of BAT, related to Figure 1.** (a-t) The proportion of cells assigned to each cluster at different housing conditions in the scRNA-seq dataset of mouse BAT-SVF.

**Supplementary Figure 2. Ambient temperature remodels intercellular crosstalk in BAT, related to Figure 2.** (a) Classification of the predicted communications in BAT based on their pattern similarity across different conditions. (b) Heat map showing the communication scores for the top ligand-receptor interactions following patterns 1, 2, and 4 across different conditions.

