## Supplementary figures and images for "Comprehensive analysis of intercellular communication in thermogenic adipose niche"

### Supplementary Figure 1

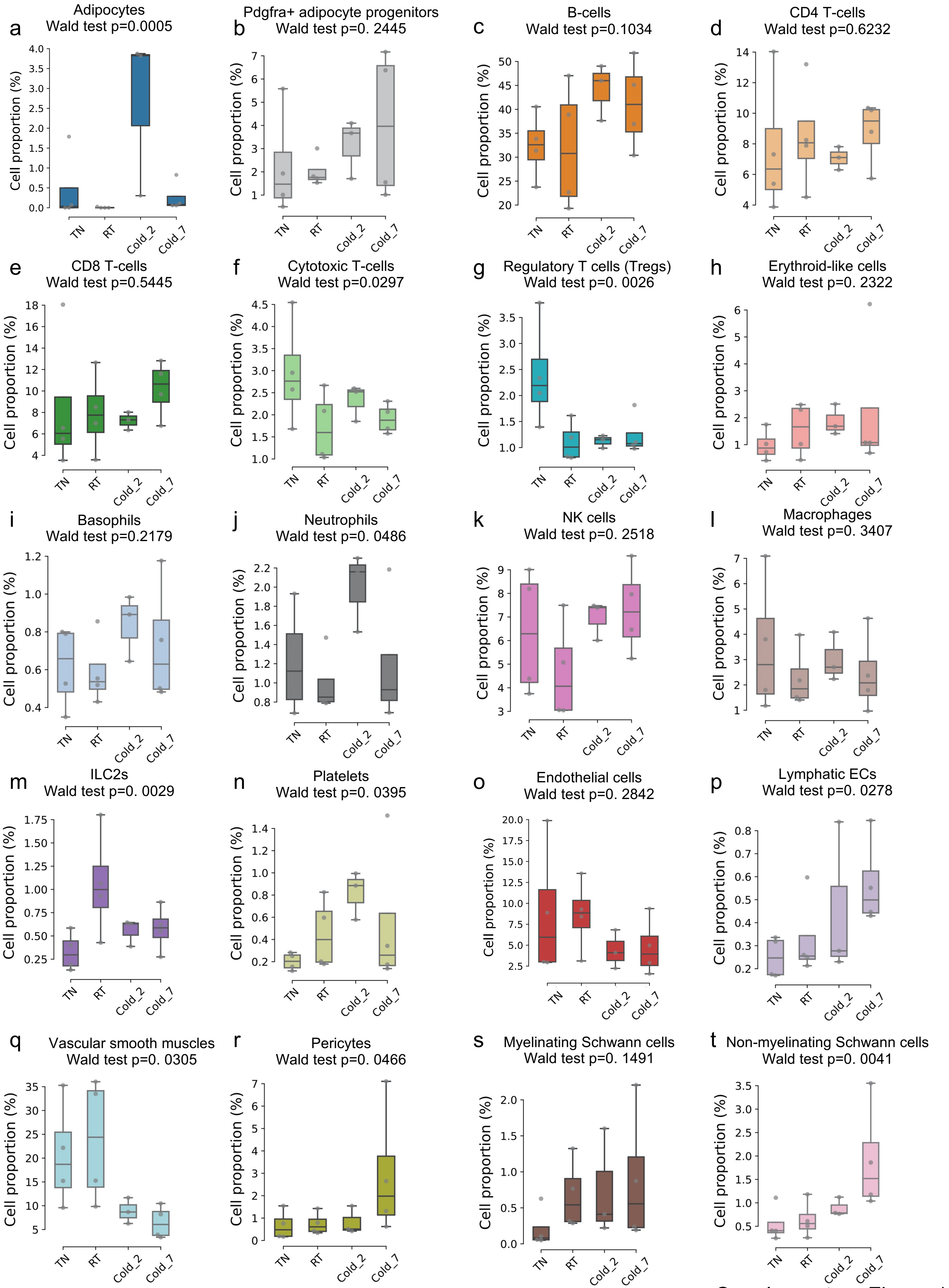


Supplementary Figure 1

### Supplementary Figure 2

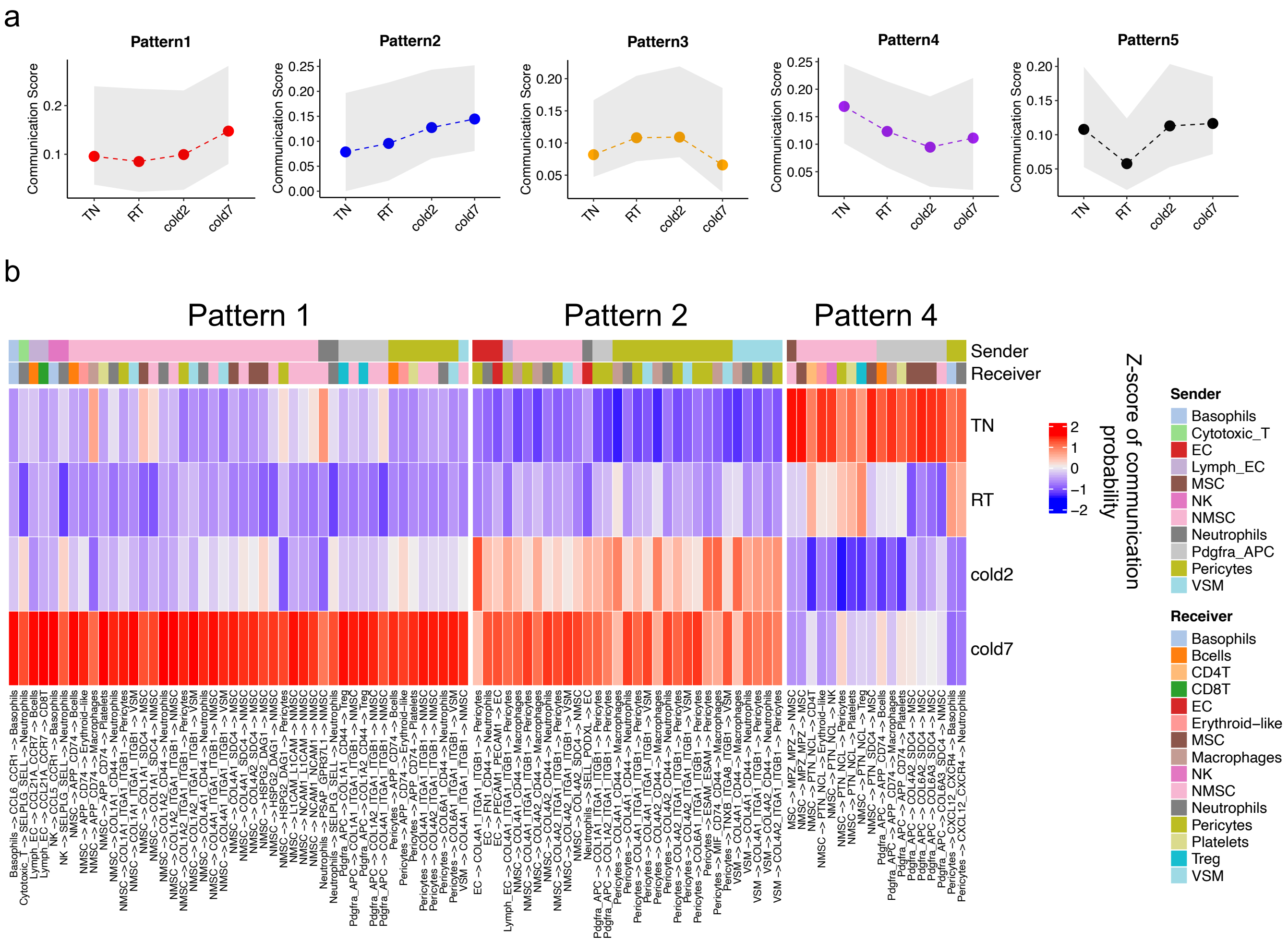


Supplementary Figure 2
